# Olfactory neuron turnover in adult *Drosophila*

**DOI:** 10.1101/2020.11.08.371096

**Authors:** Ismael Fernández-Hernández, Eric Hu, Michael A. Bonaguidi

## Abstract

Sustained neurogenesis occurs in the olfactory epithelium of several species, including humans, to support olfactory function throughout life. We recently developed a modified lineage tracing method to identify adult neurogenesis in *Drosophila*. By applying this technique, here we report on the continuous generation of Olfactory Sensory Neurons (OSN) in the antennae of adult *Drosophila*. New neurons develop sensory dendrites and project axons targeting diverse glomeruli of the antennal lobes in the brain. Furthermore, we identified sustained apoptosis of OSN in the antennae of adult flies, revealing unexpected turnover in the adult olfactory system. Our results substantiate *Drosophila* as a compelling platform to expedite research about mechanisms and compounds promoting neuronal regeneration, circuit remodeling and its contribution to behavior in the adult.

## INTRODUCTION

Sensory systems mediate the processing of external cues for the establishment of environmental and inter-individual interactions. Olfaction is essential in most species to identify food and mates, to escape from predators, and to establish memories. In order to sustain these functions throughout life, adult neurogenesis in the olfactory neuroepithelium occurs across several species, including humans (Bayramli et al., 2017; Durante et al., 2020; Fletcher et al., 2017; Graziadei and Graziadei, 1979; Graziadei and Okano, 1979; Weiler and Farbman, 1997).

Research on the olfactory system of the fruit fly *Drosophila* melanogaster has provided detailed insight on the mechanisms mediating odor coding and its evoked behaviors and memory formation at the molecular, cellular and circuitry level (Busto et al., 2010; Cognigni et al., 2018; Masse et al., 2009). Yet, generation of Olfactory Sensory Neurons (OSN) in the adult fly has not been reported thus far. Its identification would expedite a detailed analysis of the programs controlling OSN proliferation in response to environmental toxins, adult circuit remodeling, and the contribution of adult-born neurons to modulate odor-evoked behavior and memory.

The cellular architecture and function of the adult olfactory system in Drosophila is remarkably conserved to that in vertebrates, but numerically simpler. About 1300 OSN are clustered into sensory hairs called sensilla. These reside on the surface of the third antennal segment and the maxillary palps of flies, the olfactory organs equivalent to vertebrate’s olfactory epithelium. Here, most OSN express one of ~60 Olfactory Receptors (OR), as well as Orco, a related co-receptor. The axons of OSN project into diverse glomeruli in the Antennal Lobes (AL) in the brain, the counterparts to vertebrate’s olfactory bulb. Local Interneurons (LI) innervate the AL to mediate intra- and inter-glomerular inhibition. Finally, Projections Neurons (PN) convey the olfactory signals to the Mushroom Bodies (MB) and Lateral Horn (LH), the centers for integration and storage of olfactory information, equivalent to the olfactory cortex and higher processing centers in vertebrates (Galizia and Sachse, 2009; Hallem and Carlson, 2004; Masse et al., 2009; Vosshall and Laissue, 2008). Extensive genetic tools available in Drosophila grant multilevel analysis of the olfactory circuit in the adult, in a shorter time and at a lower cost than current vertebrate models.

We recently developed a modified lineage tracing system to detect generation of neurons in the optic lobes and the mechanosensory systemin the antennae of adult flies (Fernández-Hernández et al., 2019). We therefore hypothesized that the olfactory system also retains this capacity. By implementing this method, we identified sustained generation of OSN over three weeks in the antennae of adult Drosophila. New OSN develop sensory dendrites and project axons to the antennal lobes in the brain. Furthermore, we detected continuous apoptosis of OSN in the antennae, thus revealing turnover in the olfactory system of adult flies. Our results substantiate *Drosophila* as an advantageous platform to analyze and promote neuronal plasticity in the adult olfactory system at different levels.

## RESULTS

### P-MARCM detects adult-born OSN in *Drosophila*

We recently developed P-MARCM, a permanently-active lineage tracing system to label mitotically-derived cells in adult *Drosophila* tissues, including those appearing at low frequency, in a cell type-specific manner (Fernández-Hernández et al., 2019). Briefly, in this method, sustained expression of the recombinase FLP triggered by a heat shock (HS) mediates recombination of homologous chromosomes at G2 phase, allowing expression of cytoplasmic GFP and nuclear RFP in hemiclones of daughter cells upon mitosis. Specificity for labelling in the lineage is achieved by incorporation of cell type-specific GAL4 lines (Figure 1A-C). P-MARCM successfully detected adult neurogenesis and regeneration in the optic lobes and Johnston’s Organ (Fernández-Hernández et al., 2013, 2019). Because adult neurogenesis occurs in the visual and auditory/vestibular sensory systems, we hypothesized that the adult olfactory system also retains this capacity. To investigate this, we applied P-MARCM with the pan-neuronal line *nsyb-GAL4* (P-MARCM nsyb), in order to maximize the number of new OSN detected in the antennae (Vosshall et al., 2000; Sethi and Wang, 2017). We activated P-MARCM nsyb on 3-5 days-old female flies by heat shock and quantified OSN neurogenesis from confocal images of antennae dissected weekly over the next 3 weeks (Figure 1D,E). Remarkably, this approach captured OSN generated over time throughout the third antennal segment (Figure 1F). Indeed, while background labeling in non-heat-shocked flies (0 weeks, 0w) was only 4.0 +/-0.5 SEM OSN/antenna (n=27), we identified 9.7 +/- 2.7 SEM OSN/antenna at 1w (n=15); 18.4 +/-1.7 SEM OSN/antenna at 2w (n=21), and 41.0 +/-2.8 SEM OSN/antenna at 3w (n=14) (Figure 1G). Clustering antennae by the number of OSN labeled reveals continuous neurogenesis over time (Figure 1H). This is also demonstrated by an increase in the number of antennae with OSN labeled beyond background levels over time (i.e. ≥10 OSN/antenna, or mean +2 sd (4.0 + 2*2.5) = 9 OSN/antennae) (Figure 1I). Altogether, these results reveal unexpected proliferative potential in the adult *Drosophila* olfactory system.

**Figure 1.**
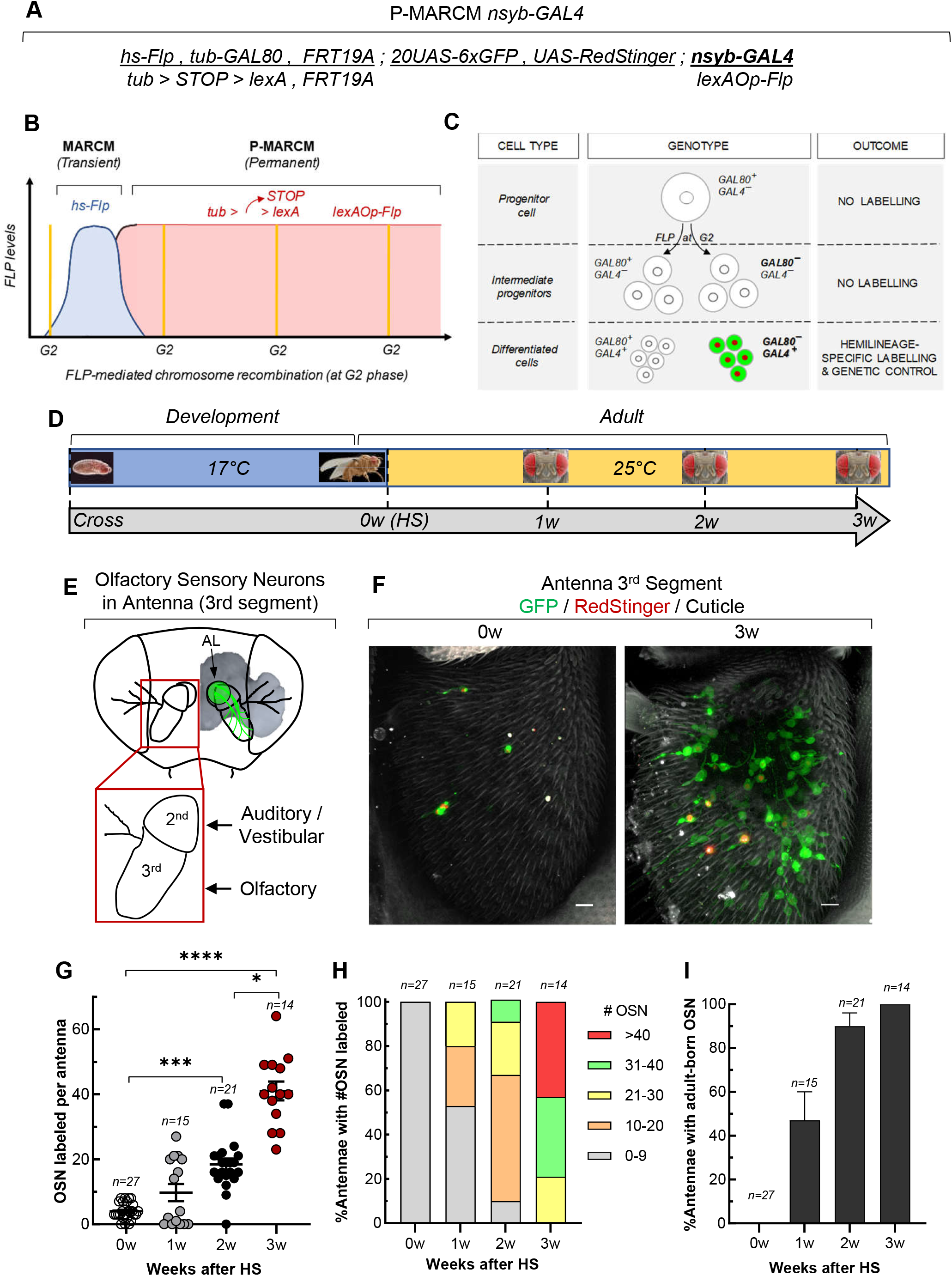
Generation of Olfactory Sensory Neurons (OSN) in the antennae of adult *Drosophila*. (A) P-MARCM is a mitotic-dependent lineage tracing system to capture adult-born neurons in *Drosophila*. (B) Sustained levels of the recombinase Flippase are expressed through the *lexA – lexA-Op* system in P-MARCM to allow mitotic recombination even in slowly dividing cells. (C) Incorporation of cell type-specific GAL4 lines into P-MARCM drives expression of cytoplasmic GFP and nuclear RFP in the cells of interest in the lineage. Additional UAS constructs allow genetic manipulation of adult-born neurons. (D) Experimental protocol to capture adult-born OSN: P-MARCM flies were kept at 17°C during development to minimize background labeling. 3-5 days-old flies were heat shocked to activate P-MARCM and antennae were dissected at 1, 2, or 3 weeks later for confocal imaging. (E) OSN located on the third antennal segment project axons through the second antennal segment to the Antennal Lobes (AL) in the brain. (F) P-MARCM reveals new OSN generated over 3w in the antennae of adult *Drosophila*. Scale bar: 10 mm. (G) Sustained generation of OSN over 3 weeks occurs in the antennae of adult flies. \**p*<0.05, \*\*\**p*<0.001, \*\*\*\**p*<0.0001. Error bars represent SEM. (H) Clustering analysis reveals continuous addition of OSN to antennae over time. (I) The number of antennae with OSN neurogenesis increases over time. By 3 weeks, 100% of the antennae have generated new OSN.

### Adult-born OSN develop dendrites and target brain circuitry

Next, we evaluated the cellular features of adult-born OSN. Firstly, detailed analysis of confocal images revealed that new OSN develop sensory dendrites, which are essential for olfaction (Figure 2A). Indeed, here the olfactory receptors are recruited to bind volatile small molecules from the environment to trigger olfaction (Larsson et al., 2004). Secondly, new OSN extend axons, which navigate through the second antennal segment towards the central brain (Figure 2B). Finally, axons of new OSN target diverse glomeruli of the Antennal Lobes (AL) in the central brain (Figure 2C). We consistently identified these features in all samples analyzed. Taken together, the cellular and circuitry features identified in new OSN strongly suggest that they mature and have the potential to functionally remodel the adult olfactory system.

**Figure 2.**
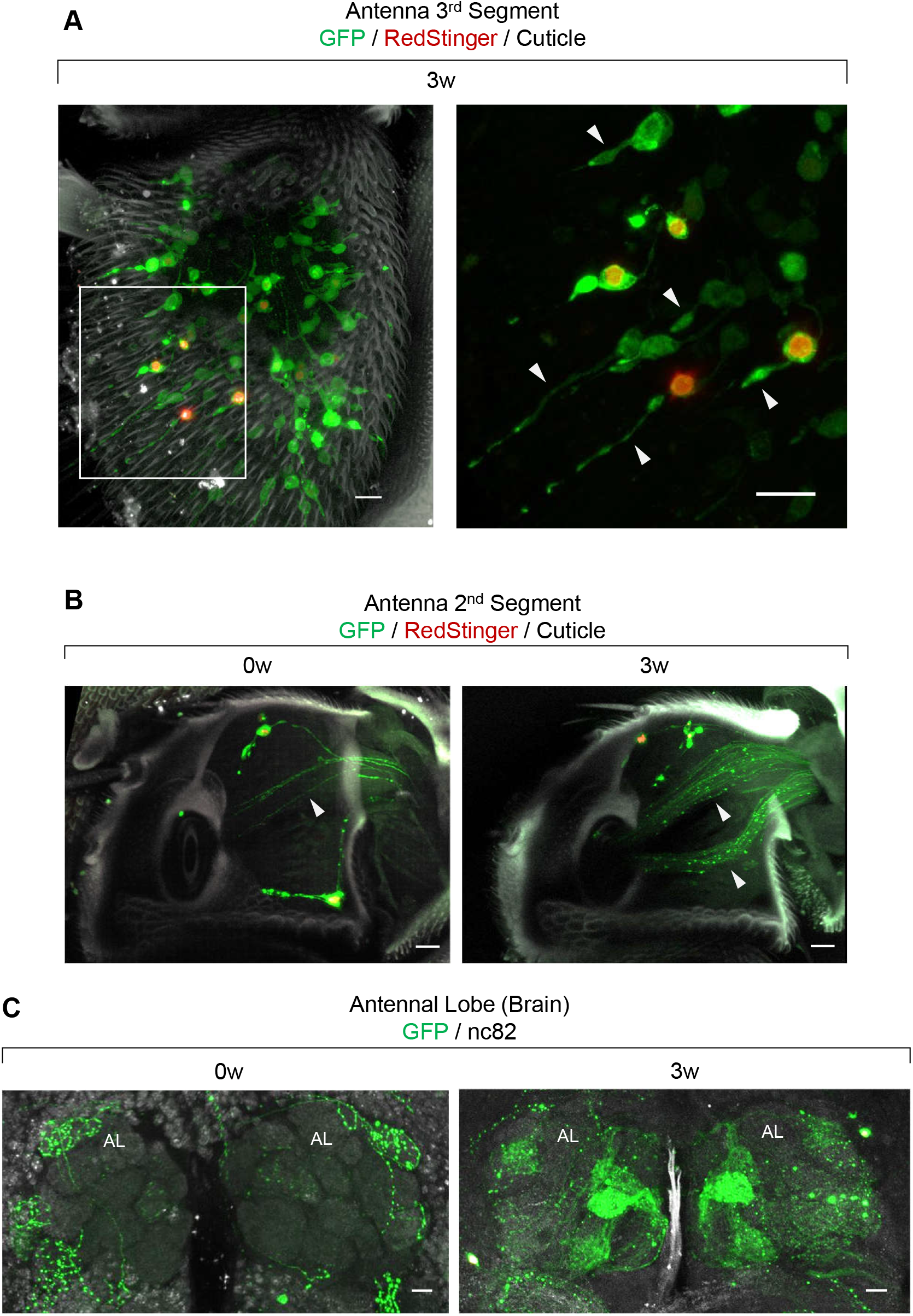
Adult-born OSN develop sensory dendrites and target brain circuitry. (A) New OSN develop sensory dendrites of different lengths projecting to sensilla. Magnification panel shows boxed area on left panel. (B) Newborn OSN in the antennae send axons to the brain through the second antennal segment (arrowheads). (C) Axons of newborn OSN target different glomeruli in the Antennal Lobes (AL) in the brain. Images presented are for the control (0w) and experimental (3w) antennae shown in Figure 1F. Scale bars for all panels: 10μm.

### Different types of OSN are generated in the adult antenna

We next evaluated whether one or more types of OSN are generated in the antenna, based on different cellular and circuit features. Firstly, the diversity of OSN in the antenna is harbored in four types of sensilla: trichoid, basiconic, intermediate, and coeloconic, which segregate into different domains on the third antennal segment (Shanbhag et al., 1999; Couto et al., 2005; Grabe et al., 2016) (Figure 3A). By applying P-MARCM, we identified new OSN throughout the third antennal segment spanning the domains of the different types of sensilla, indicating that multiple types of OSN are generated (Figure 3B). Second, OSN can also be linked to one type of sensillum by the length of their dendrites (Figure 3C). At this level, we identified OSN with long and short dendrites, characteristic of trichoid and coeloconic or small basiconic sensilla, respectively (Shanbhag et al., 1999) (Figure 3D). Third, at the circuit level, OSN expressing the same OR converge to a single glomerulus in the AL (Couto et al., 2005; Fishilevich and Vosshall, 2005; Vosshall et al., 2000) (Figure 3E). Here, we observed different glomeruli in the AL innervated by the adult-born OSN, indicating that more than one class of OSN were generated (Figure 3F). These representative features were consistently observed across the different samples analyzed. Taken together, these observations indicate that adult neurogenesis is not restricted to a single type of olfactory neurons (*i.e*. those expressing a single combination of olfactory receptors), but it rather spans to neurons of different subtypes.

**Figure 3.**
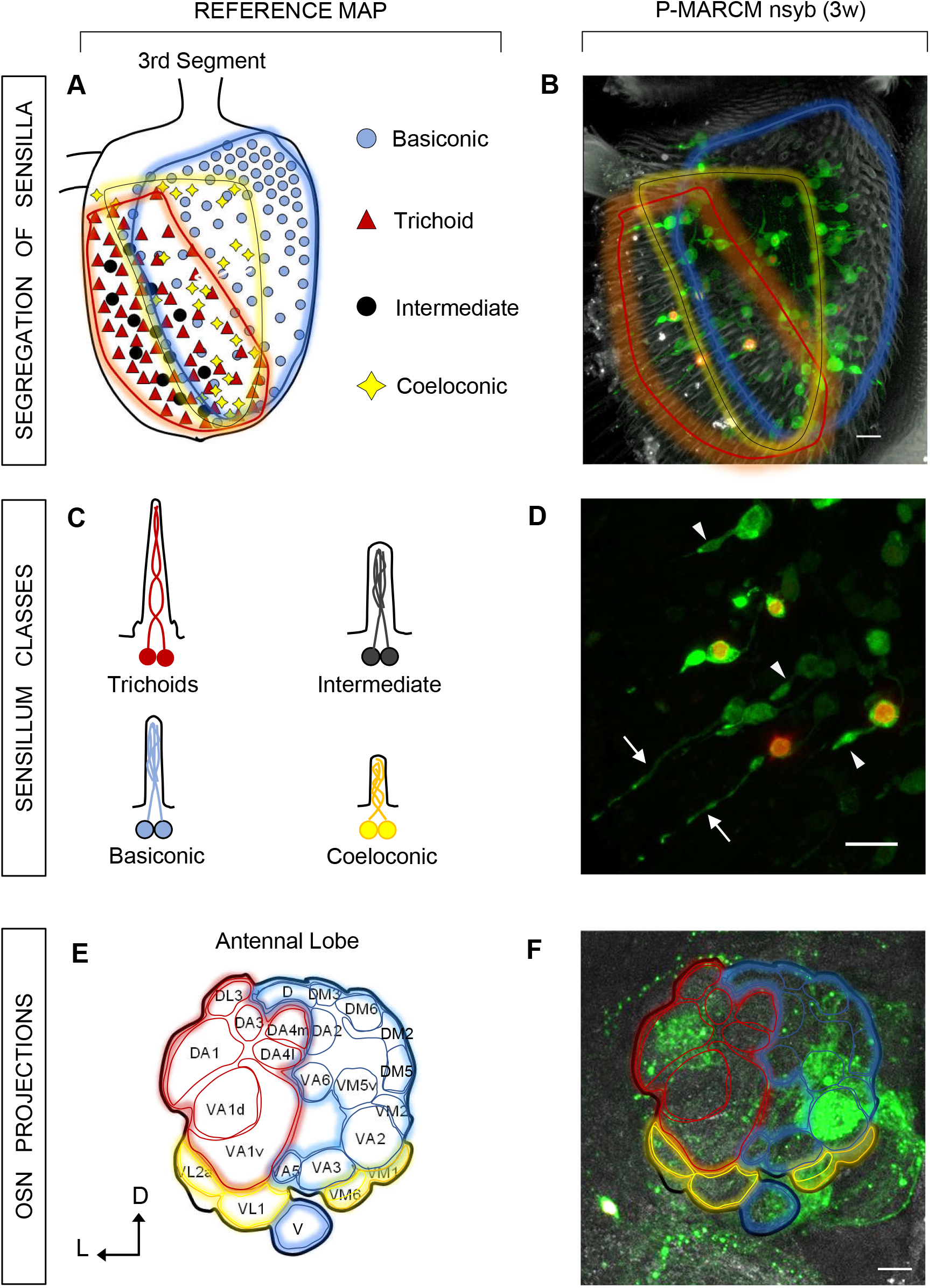
Adult neurogenesis spans to diverse OSN in the antenna. (A-B) New OSN are generated in the domains of different sensilla in the antenna. Reference map on panel A based on (Couto et al., 2005; Lin & Potter, 2015). For simplicity, the three types of Basiconic and Trichoid sensilla are represented as a single one; intermediate and Trichoid sensilla are also both surrounded by the red area. (C-D) The different lengths of sensory dendrites in newborn OSN are suggestive of neurons in the Trichoid (long, arrows) and Coeloconic (short, arrowheads) sensilla. Cartoons on panel C based on (Couto et al., 2005; Shanbhag et al., 1999). For simplicity, only sensilla containing 2 OSN are represented. (E-F) Axons of newborn OSN innervate diverse glomeruli in the AL, indicative of neurons expressing diverse olfactory receptors. Reference map on panel E shows glomeruli on the frontal view of AL, based on (Couto et al., 2005). Scale bars for all panels: 10μm.

### Turnover of adult OSN

We next asked whether new OSN detected by P-MARCM might replace lost ones in an ongoing turnover process, as it’s been suggested for the Johnston’s Organ (Fernández-Hernández et al., 2019). To answer this, we expressed the genetically-encoded reporter of apoptosis *UAS-GC3Ai* (GFP sensor Caspase-3-like protease Activity indicator (Schott et al., 2017)) with the pan-neuronal *nsyb-GAL4* driver. Briefly, expression of GC3Ai produces a non-fluorescent GFP due to an intervening caspase-recognizing sequence; upon cleavage of this sequence by active caspases, GFP becomes rapidly fluorescent in apoptotic cells (Figure 4A). By using this method, we detected GFP expression in OSN with condensed nuclei, a hallmark of apoptosis, thus validating the utility of GC3Ai as an apoptosis reporter in the *Drosophila* olfactory system (Figure 4B). Indeed, GC3Ai revealed sustained apoptosis of OSN in the antennae of female and male flies over three weeks (Figure 4C), with no significant difference between sexes at each time point analyzed (7.0+/-0.6 SEM apoptotic OSN/antenna at 1w, n=31 total antennae; 9.5+/- 0.5 SEM at 2w, n=47; 8.1+/-0.6 SEM at 3w, n=23; Figure 4D). The consistent identification of apoptotic cells at discrete timepoints strongly suggest that OSN are continuously eliminated over the analysis period in the adult.

**Figure 4.**
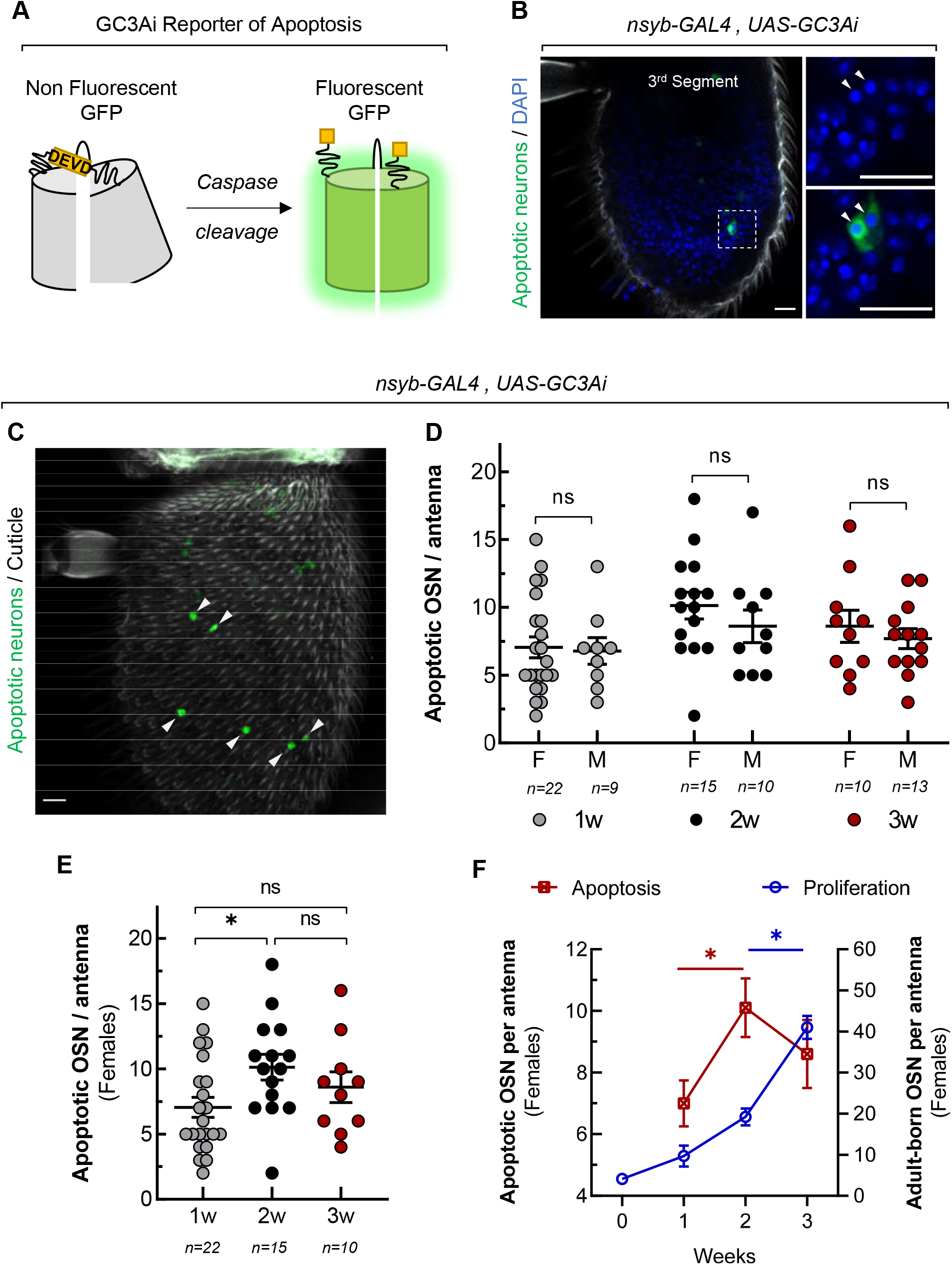
Turnover of adult OSN. (A) In the GC3Ai reporter of apoptosis (Schott et al., 2017), an inserted DEVD sequence renders GFP non fluorescent. Upon its cleavage by activated caspases, GFP reconstitutes and becomes fluorescent in apoptotic cells. (B) GC3Ai is expressed in OSN by nsyb-GAL4 line. Apoptotic OSN labeled by GC3Ai show condensed DNA (arrowheads) as a hallmark of apoptosis. (C) Apoptotic OSN are detected throughout the 3^rd^ antennal segment by *nsyb>>GC3Ai* (arrowheads). (D) Apoptosis in OSN is consistently detected in Males (M) and Females (F) with no significant difference for each time point analyzed (unpaired t-Test, *p*>0.3 for all cases). (E) A significant increase in OSN apoptosis is detected in the antennae of female flies from 1w to 2w (one-way ANOVA, * *p*=0.04). (F) Physiologic OSN turnover is revealed by a compensatory proliferation following sustained apoptosis of neurons over time. A significant increase in OSN proliferation at 3w following a significant increase in OSN apoptosis at 2w further substantiates this interpretation. Scale bars: 10μm. Error bars represent SEM.

In order to correlate OSN apoptosis with proliferation, we focused on data from female flies, since P-MARCM relies on X chromosome mitotic recombination and labelling, and is therefore active only in females. Statistical analysis reveals a significant increase in OSN apoptosis in 2w-old female flies (10.1 +/-1.0 SEM OSN/antenna, n=15) with respect to 1w-old flies (7.0 +/-0.8 SEM OSN/antenna, n=22; *p*=0.04, one-way ANOVA, Figure 4E). This correlates with a compensatory, significant increase in OSN proliferation from 2w (19.2 +/-2.1 SEM OSN/antenna, n=17) to 3w (41.0 +/-2.8 SEM OSN/antenna, n=14; *p*=0.04, Kruskal-Wallis test, Figure 4F). Taken together, the sustained levels of apoptosis detected by GC3Ai and the continuous proliferation captured by P-MARCM over three weeks indicate turnover of OSN, revealing unexpected cellular plasticity in the adult *Drosophila* olfactory system.

### Adult neurogenesis across sensory systems in *Drosophila*

Finally, we compared adult neurogenesis in the olfactory system with that occurring in other sensory systems. New neurons have also been detected by P-MARCM in the visual and auditory/vestibular system of 3 week-old adult flies (Fernández-Hernández et al., 2019) (Supplementary Figure 1A-C). A comparative analysis across the 3 sensory systems reveals the highest rate of neurogenesis in the olfactory system of the third antennal segment (41.0+/-2.8 SEM OSN/antenna, ~3% of ~1300 total OSN per antenna,100% antennae (n=14), this work), followed by the mechanosensory auditory/vestibular JO neurons in the second antennal segment (11.2 JO neurons/antenna, ~2.2% of ~500 JO neurons per antenna, interpolation for 3 weeks from 2 weeks (n=7) and 4 weeks (n=7) timepoints, 50% of antennae analyzed), and lastly by the projection neurons in the medulla of the optic lobes 31.0+/- 2.4 SEM neurons per optic lobe, ~0.1% of ~40,000 interneurons, 100% optic lobes analyzed (n=12), (Fernández-Hernández et al., 2019) (Supplementary Figure 1D). These differences in proliferation may reflect different levels of cell death on each system. Indeed, while projection neurons in the optic lobes seem to be much more protected from external damage, the OSN in the antennae are permanently exposed to potential environmental insults which might ultimately lead to their elimination. Yet, the neurogenesis identified reflects a regenerative capacity evolved on each sensory system to repair and potentially restore their functions.

## DISCUSSION

Cellular plasticity occurs in defined regions of the adult nervous system. Continuous neurogenesis in the olfactory neuroepithelium supports olfactory function in different species. By implementing the P-MARCM lineage tracing method with the pan-neuronal *nsby-GAL4* line, here we report sustained generation of OSN over 3 weeks in the antennae of adult *Drosophila* (Figure 1). This is revealed by an increase in i) the number of OSN generated per antenna, and ii) the number of antennae undergoing OSN neurogenesis over time. We identified adult neurogenesis in 100% of the antennae analyzed at 3 weeks (n=14), with a mean of 41.0 +/-2.8 SEM new OSN/antenna, which represents ~3% turnover of the ~1300 OSN in each antenna. Importantly, because P-MARCM labels only hemi-clones from cells in which the system gets activated upon heat-shock, the actual amount of new OSN might be higher. Furthermore, a detailed analysis of P-MARCM images revealed that new OSN develop sensory dendrites and project axons to the AL (Figure 2). These cellular features strongly suggest a complete maturation and integration of new OSN into pre-existing circuitry.

Three lines of evidence suggest that more than one type of OSN are generated in the antennae, by comparing cellular features of new OSN against reference maps (Vosshall et al., 2000; Couto et al., 2005; Grabe et al., 2016): (i) at the topological level, new OSN are scattered throughout the domains of the different sensilla in the third antennal segment; (ii) at the cellular level, new OSN develop dendrites of different lengths: long for Trichoids and short for Coeloconic or small Basiconinc; (iii) at circuitry level, axons of new OSN target different glomeruli in the antennal lobes (Figure 3). An unambiguous assessment of the OSN subtypes generated can be achieved by replacing *nsby-GAL4* in P-MARCM by Olfactory Receptor-specific *GAL4* drivers (Vosshall et al., 2000; Couto et al., 2005; Fishilevich and Vosshall, 2005; Lin and Potter, 2015). Yet, our results indicate that neurogenesis spans to different OSN types in the antennae of adult flies over 3 weeks.

We also identified sustained apoptosis of OSN in physiologic conditions, revealed by the genetically-encoded GC3Ai reporter (Figure 4). Indeed, apoptotic OSN were captured in the antennae of male and female flies over 3 weeks at comparable extents. This cell death might be a consequence of ageing or the accumulation of eventual insults from the environment. *Drosophila*, as many other animals, strongly rely on olfaction for critical responses and behaviors, such as selection of food, escape from predators and selection of partners for mating. Thus, a compensatory proliferation following elimination of OSN should be in place to preserve these critical functions. Our results showing apoptosis and generation of OSN over time support an ongoing turnover in the olfactory system of adult flies. Whether OSN regenerate following injury, as it occurs in vertebrates (Graziadei et al., 1978; McM Carr and Farbman, 1992; Schwob et al., 1995; Frontera et al., 2016; Cervino et al., 2017), and how ageing affects this capacity, are questions that can be addressed in future experiments using the platform presented here. Furthermore, this platform will facilitate and expedite the screening of molecules promoting proliferation and integration of neurons in an entire adult circuit (Fernández-Hernández et al., 2016).

While P-MARCM specifically labels adult-born OSN, it does not identify their cell-of-origin. Low-rate self-division of mechanosensory neurons has been recently proposed as a mechanisms driving regeneration in the adult Johnston’s Organ (Fernández-Hernández et al., 2019). Whether this mechanism also operates in the olfactory system, or other cell types act as progenitors, can be addressed by combining cell type-specific lineage tracing systems with transcriptomic analysis (Fletcher et al., 2017; Li et al., 2020).

What is the functional role of new OSN? In vertebrates, adult-born olfactory neurons mature and innervate the olfactory bulb to support olfaction (Schwob, 2002)(Blanco-Hernández et al., 2012)(Hurtt et al., 1988)(Ducray et al., 2002). Although we did not test the function of new OSN here, we observed that they develop sensory dendrites and innervate diverse glomeruli in the antennal lobes in the brain, strongly suggesting that they mature, integrate, and remodel the olfactory circuit to sustain its function. Inhibition of programmed cell death in otherwise eliminated OSN during development allows their functional integration in the adult (Prieto-Godino et al., 2020), demonstrating a high degree of plasticity in the olfactory circuit. This capacity further supports the notion that adult-born OSN might be functionally integrated to modulate olfaction. Furthermore, the apparent diversity of OSN generated suggest their potential to respond to multiple odorant molecules, and ultimately to modulate olfactory-related behavior and memory more precisely. These questions can be approached by leveraging the versatility of P-MARCM with the multiple genetic tools available in *Drosophila*. For instance, the functional integration of newborn OSN can be assessed by circuitry-mapping of axons innervating the AL (Laissue et al., 1999)(Couto et al., 2005)(Grabe et al., 2016), in conjunction with Trans-TANGO (Talay et al., 2017), nsyb-GRASP (Macpherson et al., 2015), and calcium indicators (Chen et al., 2013; Dana et al., 2019) tools. Similarly, their behavioral contribution can be tested by applying established protocols upon selective activation of adult-born OSN by optogenetic tools (Klapoetke et al., 2014), or their genetic silencing, either in a constitutive manner, by expressing TeTx (Sweeney et al., 1995), or in a reversible manner, by expression of shi^ts^ (Kitamoto, 2001).

In summary, our results demonstrate unexpected plasticity in the olfactory system of adult *Drosophila,* and establish a unique platform to expedite the identification of molecular mechanisms and the *in vivo* screening for compounds promoting proliferation, circuit integration and the functional contribution of new neurons in the adult.

## MATERIALS AND METHODS

### Fly lines and experimental conditions

#### For P-MARCM-nsyb experiments

The P-MARCM system has been previously described (Fernández-Hernández et al., 2019). Flies of final genotype *hs-Flp,tub-GAL80,neoFRT19A / tub FRT STOP FRT lexA,neoFRT19A; 20UAS-6GFPmyr, UAS-RedStinger / +; nsyb-GAL4 / 8lexAOp-Flp* were used to assess proliferation of OSN. The parental cross was set and kept at 17°C during development to minimize FLP induction. Female flies 2-5 days old were picked and blindly assigned to either control group for dissection, or to experimental groups for heat shock at 38°C (Ohlstein and Spradling, 2006) for 45 minutes, twice on the same day ~2 hours apart. Flies were then kept at 25°C. Antennae and brains were dissected weekly afterwards over 3 weeks.

#### For GC3Ai experiments

We crossed female virgins of the apoptotic reporter *UAS-GC3Ai* (Schott et al., 2017) to *nsyb-GAL4* males. Parental cross and progeny were kept at 25°C. Antennae of males and females of final genotype *w; +; nsyb-GAL4 / UAS-GC3Ai* were dissected at 1, 2 and 3 weeks after eclossion for confocal imaging as described below.

#### Dissection and immunostaining

Antennae were dissected, attached to their corresponding brains in chilled Schneider’s medium and then fixed in 3.7% formaldehyde solution for 20 min. They were then washed with PBS + Triton (1%) solution for 20 min, followed by a final wash in 1XPBS before incubation with primary nc-82 antibody overnight at 4°C, followed by incubation with secondary antibody 4 hours at room temperature or overnight at 4°C. Antibodies used were mouse anti-nc82 (1:10, DSHB) and secondary antibody anti-mouse Cy5 (1:100, Jackson laboratories). No antibodies were used for GFP and RedStinger fluorescent proteins. Antennae and brains were mounted in Vectashield media with DAPI (Vector laboratories). For mounting, we used double-side sticker spacers (EMS, 70327-9S) to preserve morphology as much as possible (one for antennae, two for brains).

#### Statistical analysis

Plots and statistical analysis were done using GraphPad Prism, applying Kruskal-Wallis test for multiple comparisons in Figure 1G and 4F, unpaired t-Test for data on Figure 4D and one-way ANOVA for data on Figure 4E. Error bars represent SEM in all plots. SEM on experiments with binary outcomes (Figure 1I) was calculated as 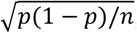, where p is the frequency of OSN neurogenesis in the analyzed group and n the number of flies considered. For clustering of antennae based on the number of OSN labeled (Figure 1H), any antennae with a given number of OSN higher than the mean +2 sd in the control non-heat-shocked group (i.e. background) was regarded as adult neurogenesis. This yielded a threshold of 9.0 OSN/antenna, thus establishing the first cluster as those antennae with 0-9 OSN neurons as background and any values above this as adult neurogenesis.

## ACKNOWLEDGEMENTS

We thank members of the Bonaguidi Lab for support; Bloomington Drosophila Stock Center (NIH P40OD018537) for *UAS-GC3Ai* and *nsyb-GAL4* lines used in this study; and Developmental Studies Hybridoma Bank, created by the NICHD of the NIH for nc-82 antibodies.

## AUTHOR’S CONTRIBUTIONS STATEMENT

IFH designed and conducted experiments and carried out data collection and analysis. EH conducted experiments. MB supervised experiments, provided funding and resources, and provided input for the manuscript. IFH wrote the manuscript with input from all authors.

---

**Supplementary Figure 1.**
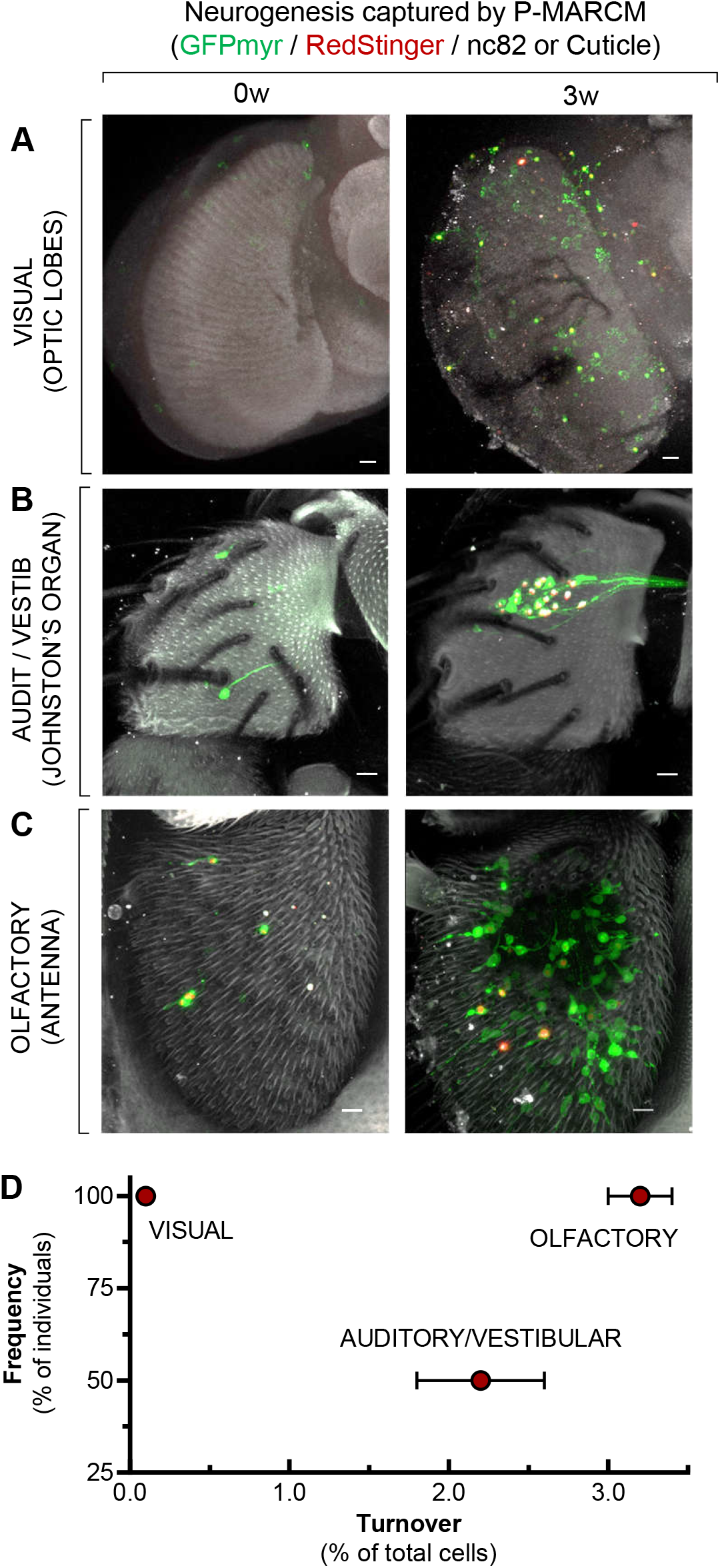
Adult neurogenesis accross sensory systems in *Drosophila*. (A-C) P-MARCM captures neurogenesis over 3 weeks in the Visual system (Optic Lobes) (A), the Auditory/Vestibular system (JO) (B), and the Olfactory System (antena) (C) of adult *Drosophila*. For JO, a representative image of neurogenesis at 4w is shown. Scale bars: 10 μm. (D) The Olfactory system shows the highest neurogenesis turnover at 3w among the three sensory systems analyzed. For auditory/vestibular, an interpolation of 2w and 4w data is plotted (Fernández-Hernández et al., 2019). Error bars represent SEM.

